# Distinctive fragmentation of circulating DNA from chromosome Y and its alteration in cancer

**DOI:** 10.64898/2026.11.22.768410

**Authors:** Léa Ruffinel, Marianne Richaud, Ekaterina Pisareva, Alain R. Thierry, Jacques Colinge

## Abstract

Circulating cell-free DNA (cirDNA) fragmentation reflects chromatin organization and represents a promising source of cancer biomarkers. Here, we characterized the fragmentation patterns and nucleosome occupancy of Y chromosome (chrY) cirDNA in healthy individuals and cancer patients. In healthy males, we found that the shorter chrY-derived cirDNA fragments predominantly originate from heterochromatic regions. We also observed substantially lower nucleosome occupancy on chrY than on all other chromosomes, a finding independently supported by micrococcal nuclease digestion data. Remarkably, in patients with colorectal, pancreatic, lung and prostate cancer, the chrY-specific enrichment of shorter fragments was markedly attenuated, contrasting with the increased fragmentation usually observed in cancer when chromosome specificity is disregarded. We reproduced these observations in an independent local cohort of healthy and colorectal cancer patients. We could exploit ratios of short (100–147 bp) to long (147–200 bp) fragments between chrY and other chromosomes to discriminate healthy males from patients with colorectal, pancreatic, and lung cancers with AUCs above 0.98. Autosomes 19, 17, and 22 also displayed atypical fragment patterns, different from chrY and associated with gene density. This last observation was confirmed in females.

## INTRODUCTION

Cell-free circulating DNA (cirDNA) collected in blood originates from tissues all over the body, but hematopoietic cells constitute the main source (1). CirDNA is released through several cellular processes (apoptosis, necrosis, NETosis, etc.). Upon release, it is fragmented by multiple enzymes (DNAses) in the cytoplasm first, and in blood circulation subsequently (2). Importantly, DNA bound to proteins such as histones is partially protected from DNAses (3). This results in a cirDNA fragment length profile featuring a characteristic major mode at 167 bp, which corresponds to the length of DNA wrapped around a nucleosome with linker histone H1, *i.e.*, a chromatosome (4). The high reproducibility of this length distribution among healthy individuals and its alteration in certain diseases, *e.g.*, cancer, make it a source of systemic biomarkers. Other characteristics of cirDNA fragments such as their end sequences (end motifs) display reproducible variation between healthy and disease, thereby enriching the repertoire of cirDNA biomarkers (5–9). Moreover, due to the aforementioned protective effect of histones against DNAses, multiple studies have shown that cirDNA can be used to infer nucleosome positions and occupancy (10,11).

The Y chromosome (chrY) stands out among human chromosomes. It is the shortest chromosome, spanning approximately 57-62 million base pairs (bp) and only encoding 78-106 protein-coding genes (12,13). It contains a male-specific region along with two pseudo-autosomal regions (PAR), which share homology with regions of the X chromosome (chrX), and a large heterochromatin region enriched in repeated sequences (13,14). Another notable characteristic of chrY is mosaic loss, a phenomenon particularly associated with aging where certain somatic cells lose chrY (15). ChrY plays a critical role in male development, fertility, and sex-linked disorders. Its loss or alterations have been associated with increased cancer risk, immune dysfunction, and aging-related pathologies (16).

Despite the increasing application of cirDNA in a number of research projects, the chromosome-specific fragmentation properties of cirDNA remain understudied. Recently, we discovered that chrY presents shorter fragments compared to those originating from other chromosomes (17). In this study, we refined this initial observation by characterizing more precisely the origin of the shorter fragments, how their abundance changes in cancer, and chrY nucleosome occupancy. To put this work in a broader context, we also explored the possibility that cirDNA originating from other chromosomes would harbor distinctive fragmentation and nucleosome occupancy patterns.

## RESULTS

### ChrY Fragment length distribution in healthy individuals

We used the healthy cohort from Cristiano, *et al*. (5) that comprises 81 male samples out of 245, excluding technical replicates. CirDNA fragments from males only were used. Considering the autosomes and chrX without the PAR (auto+X) on the one hand, and chrY without the PAR on the other, we reproduced our previous observation made on an independent cohort (17). ChrY fragments were enriched in smaller lengths (**Fig. 1A**). To better represent this bias, we adopted a representation of the fragment length distribution differences chrY *versus* auto+X. Fragments shorter than 147 bp were underrepresented in auto+X relative to chrY, whereas fragments between 147 and 200 bp were overrepresented (**Fig. 1B**). The transition at 147 bp corresponds to the length of a mononucleosome. The most overrepresented fragment length in auto+X was 167 bp, corresponding to the length of a mononucleosome with histone H1.

**Figure 1.**
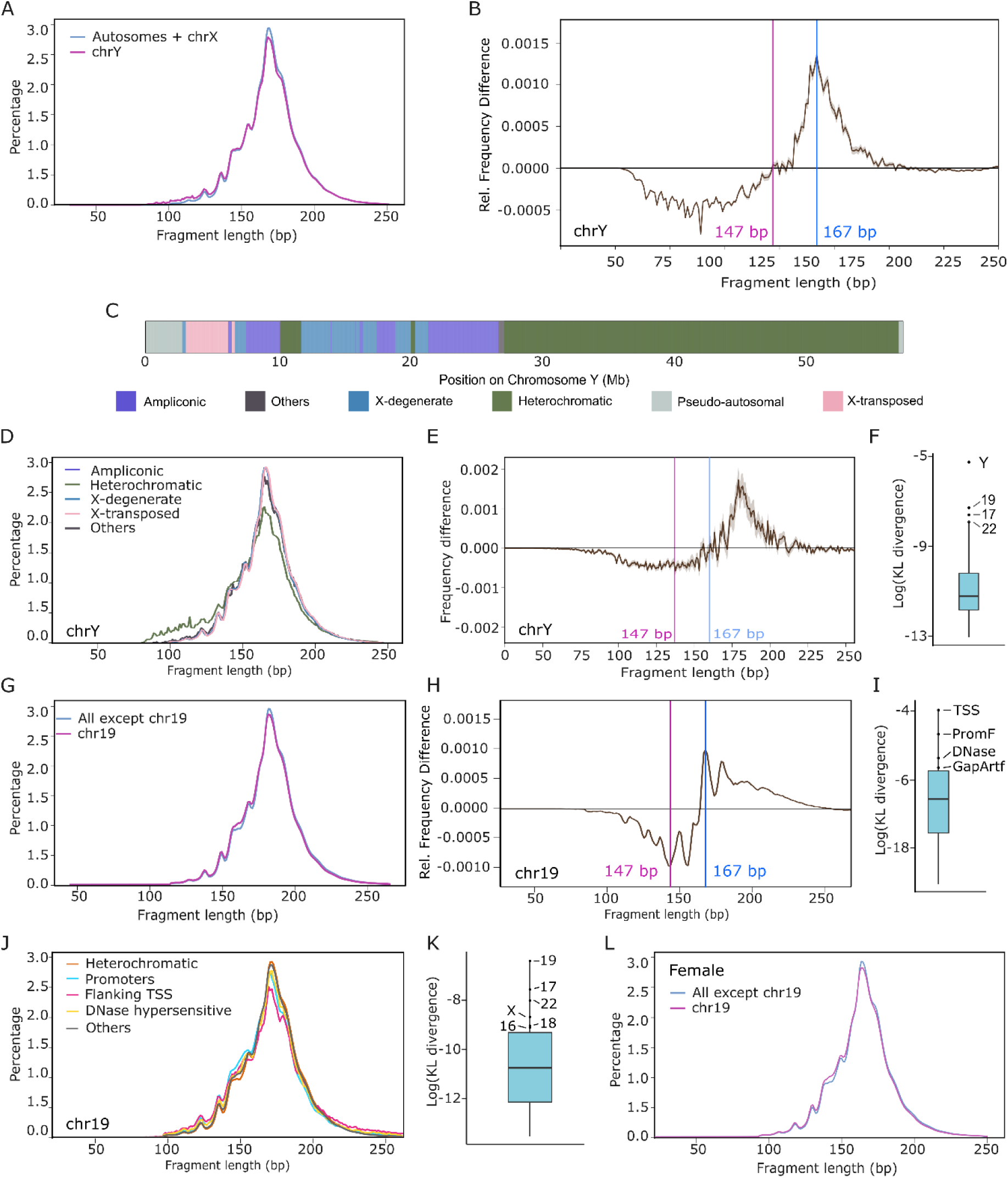
(**A**) CirDNA fragment length distributions of male healthy individuals (Cristiano dataset) for fragments mapped on chrY *versus* autosomes and chrX (auto+X). (PAR were excluded in both cases.) (**B**) Fragment length relative frequency differences between auto+X and chrY. (**C**) Annotations of chrY subregions. (**D**) Fragment length distributions of male healthy individuals mapped on chrY for each subregion type. (**E**) Fragment length relative frequency differences between auto+X and chrY for the local healthy dataset. (**F**) Kullback-Leibler divergence (KLD) of the fragment length distribution of each chromosome *versus* all the others (male individuals only). Chromosomes 19, 17, and 22 also appear as outliers. (**G**) Chr19 cirDNA fragment length distribution versus all the others (males only). (**H**) Fragment length relative frequency differences between chr19 and all the others. (**I**) KLD of fragment length distributions associated with genomic annotations (TSS = transcription start site, PromF = flanking promoter, DNase = DNAse hypersensitive, GapArtf = alignment artifact). (**J**) Chr19 fragment length distributions *per* selected genomic region annotation type. (**K**) KLD of the fragment length distribution of each chromosome *versus* all the others (female individuals only). (**L**) Chr19 cirDNA fragment length distribution versus all the others (females only).

To further dissect the fragmentation of chrY cirDNA, we used the subregion annotations proposed by Hallast, *et al*. (14) (**Fig. 1C**). For each annotated category, we computed the Kullback-Leibler divergence (KLD) between the length distributions of fragments from the particular category and those outside the category. The heterochromatic regions were the most enriched in shorter cirDNA fragments (KLD = 0.078, other categories KLD < 0.012) (**Fig. 1D**).

We validated our observations on a local healthy cohort comprising 20 male samples (**Fig. 1E, S1A**). In this cohort, we observed that fragments shorter than 167 bp were underrepresented in auto+X relative to chrY, whereas fragments between 167 and 210 bp were overrepresented. Additionally, the enrichment in short fragments from heterochromatic regions was reproducible on the local cohort (**Fig. S1B**).

### Atypical fragment length distributions in other chromosomes

To complement our observations on chrY, we considered all the autosomes and chrX. We computed each chromosome specific fragment length distribution and compared it to the distribution of all the other chromosomes pooled. We performed this analysis using male samples only to have the same population as with the chrY. The KLD was applied to characterize the divergence of a particular chromosome, which revealed that chrY, chromosome 19 (chr19), chr17, and chr22 displayed clear differences (**Fig. 1F**). The strongest divergence after chrY was from chr19 (**Figs. 1GH**), the plots for chr17, chr22, and all the nondivergent chromosomes are in **Figures S2 & S3**. Fragment length distributions for chr19, chr17 and chr22 in the local healthy cohort also displayed slight divergence when compared to all other chromosomes (**Fig. S4**).

Interestingly, chrY divergence was primarily located outside the 147-167 bp interval, while for chr19, chr17, and chr22, the divergence was primarily within this interval. Since the latter chromosomes count among the gene denser ones, we thought that this atypical pattern might relate to gene expression and, in particular, transcription regulatory regions such as transcription start sites (TSS) and promoters (18,19). We hence correlated fragment length with genomic regions as annotated in Vu *et al.* (20). We identified TSS, flanking promoters, and DNAse hypersensitive regions being the most biased according to KLD (**Fig. 1IJ**). The GapArtf corresponds to alignment artifacts and was thus not biologically relevant. For comparison, heterochromatic regions were slightly enriched in larger fragments (**Fig. 1J**). The same associations held true for chr17 and chr22 (**Fig. S5**). To be complete, autosome and chrX analyses were also performed using Cristiano 164 healthy female samples. Chr19, chr17, and chr22 were also the most divergent, followed by chrX. Chr18 and chr16 were marginally divergent (**Figs. 1KL, S6**).

### Nucleosome occupancy

We found a stable number of cirDNA fragments *per* chromosome across male healthy individuals in autosomes after normalizing for their respective lengths (**Fig. 2A**). A slight decrease in fragment number was observed for chromosomes 9, 13, 14, 15, 21, and 22. Those chromosomes contained large regions with almost no detected cirDNA fragments. These regions were excluded from our study. They correspond in majority to acrocentric chromosome short arms (21). The number of fragments from chrX was approximately half that of the autosomes, consistent with its haploid state in males. However, chrY was associated with a considerably lower number of fragments even compared to chrX. We hypothesized that this was due to lower nucleosome occupancy, resulting in augmented degradation of fragments less protected by histones.

**Figure 2.**
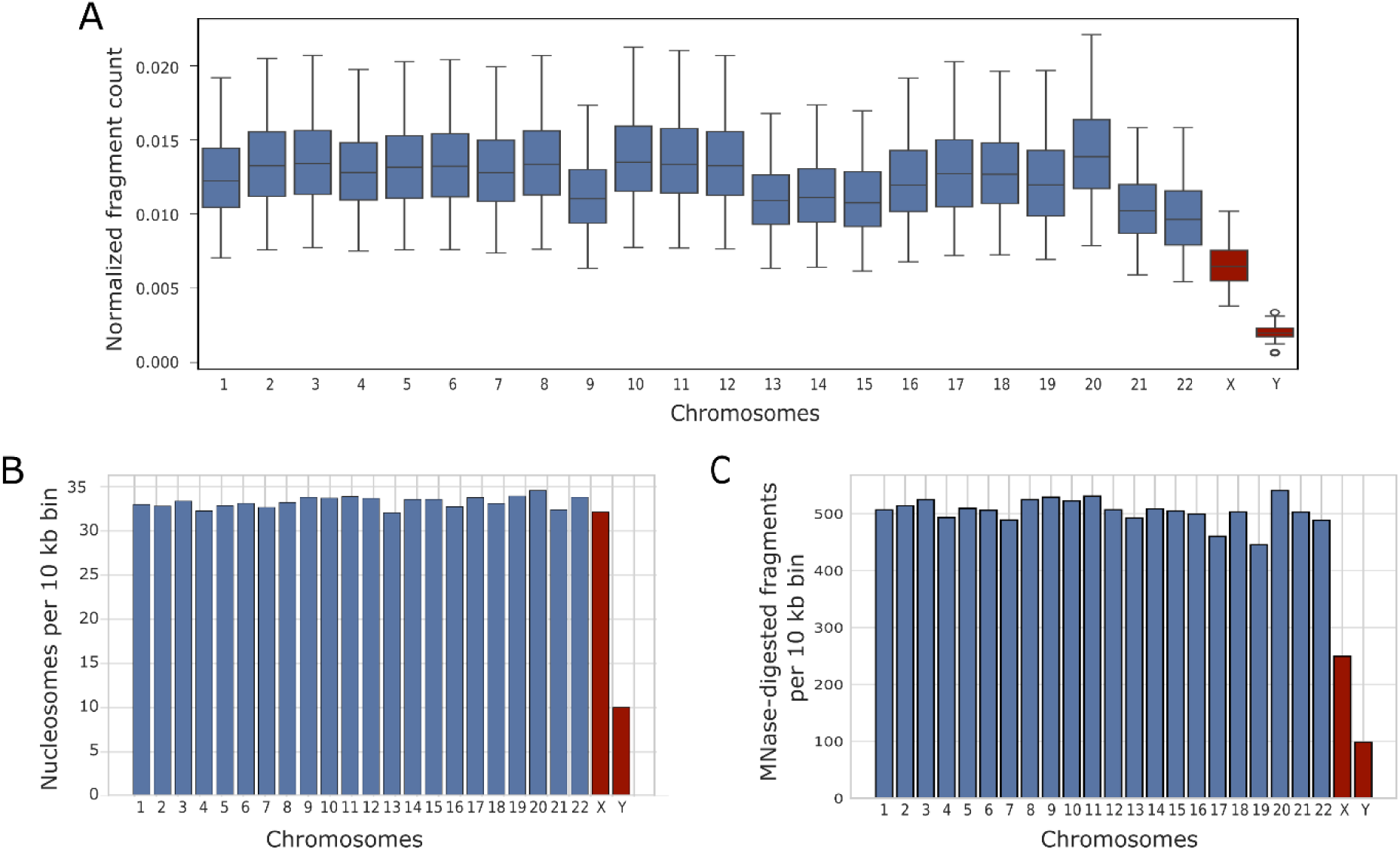
**(A**) Number of fragments mapped on each chromosome normalized by chromosome size for healthy male individuals. (**B**) Number of stable nucleosomes *per* 10 kb bin on each chromosome for healthy male individuals. (**C**) Number of MNase-digested fragments per 10 kb bin on each chromosome for CD4+ T cells (Schones, *et al.* data).

To test this hypothesis, we inferred stable nucleosome positions using the total cirDNA genome coverage of male samples. For this purpose, we used an algorithm that we recently published, which identifies cirDNA coverage peaks of a diameter compatible with a nucleosome (11). The number of detected nucleosomes *per* chromosome was highly stable across autosomes and chrX, but considerably lower for chrY (**Fig. 2B**). This phenomenon was rather uniform over chrY and, in particular, no clear association with the regions in **Figure 1C** was found.

Despite the reduced number of cirDNA fragments originating from chrX by a factor 0.5, we found a density of nucleosomes on chrX comparable to the diploid autosomes. This indicated that our nucleosome detection was not perturbed by the lower chrX coverage. Nevertheless, since the number of fragments from chrY was further reduced (**Fig. 2A**), the possibility remained that the much lower nucleosome density on chrY might be due to a failure of our algorithm working with even lower chromosome coverage. We hence sought to validate our results in **Figure 2B** with an independent data set. We used micrococcal nuclease (MNase)-digested fragments corresponding to nucleosome cores (22) from a CD4+ T-lymphocyte cell line (23). Since cirDNA originates predominantly from hematopoietic cells (1,24), we considered this cell line an acceptable approximation of cirDNA composition. The number of MNase-digested fragments was stable across autosomes (**Fig. 2C**). Consistent with the haploid state of chrX in males, the number of fragments on chrX corresponded to approximately half that of the autosomes (cell line derived from a male individual as assessed by fragments mapped on chrY outside the PAR). However, the number of MNase-digested fragments on chrY was considerably lower than on chrX, thereby confirming our estimates.

### Variation in cancer

To assess whether the number of detected nucleosomes was altered in cancer, we needed a large cohort of male cancer patients. Indeed, our nucleosome detection algorithm relies on the total, population-wide genome coverage at every bp, and Cristiano individual cancer type cohorts were too limited in male samples. We hence exploited the much larger Adalsteinsson prostate cancer cohort (557 patients) (25). Results highly similar to the healthy case were obtained (**Fig. 3A**).

**Figure 3.**
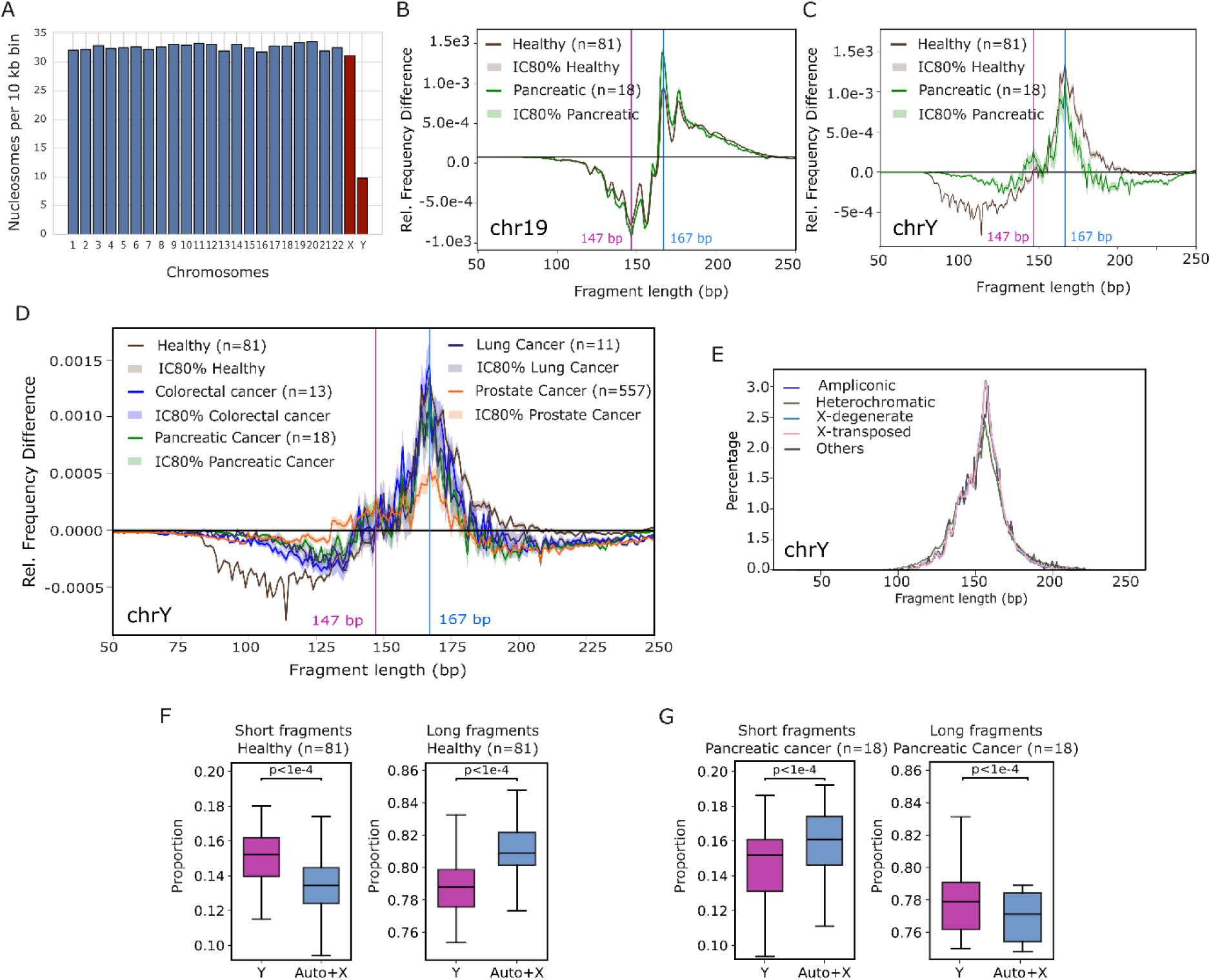
(**A**) Number of stable nucleosomes per 10 kb bin on each chromosome for prostate cancer patients (Adalsteinsson cohort). (**B**) Fragment length relative frequency differences between chr19 and all the other chromosomes in male healthy individuals and pancreatic cancer patients (Cristiano cohorts). (**C**) Same analysis for chr1Y against auto+X. (**D**) Fragment length relative frequency differences between auto+X and chrY fragments in male healthy, colorectal, lung and pancreatic cancer patients (Cristiano cohorts) as well as prostate cancer patients (Adalsteinsson cohort). (**E**) Fragment length distributions of male colorectal cancer patients mapped on chrY for each subregion type. (**F**) Boxplots of short and long fragments proportions in male healthy individuals (Cristiano cohort). (**G**) Boxplots of short and long fragments proportions in male colorectal cancer patients (Cristiano cohort).

We next considered the influence of cancer on fragment length distributions. This analysis was compatible with the number of males in Cristiano cancer cohorts. We considered the cancers with most males: colorectal (13 males out of 27 samples), lung (11 out of the 40 preoperative and pretreatment samples), pancreas (18 out of 35), and the prostate cancer cohort from Adalsteinsson et al. (25). Length distributions were not really distinguishable for chr19 as illustrated for pancreatic cancer in **Figure 3B**, neither was it for chr17 nor chr22 (**Fig. S7**). The observed bias found in healthy individuals remained unperturbed by cancer.

For chrY, on the contrary, we found cancer-induced variation in the length distributions as illustrated for pancreatic cancer in **Figure 3C**. Both the depletion of fragments below 147 bp and the overrepresentation of fragments between 147 and 200 bp were much less pronounced compared to the healthy condition. This observation was reproducible in the other cancer types we considered (**Fig. 3D**), which was surprising since it is well-known that in many cancer types, fragment lengths are enriched in shorter values and the Cristiano cohorts followed this general trend with all chromosomes pooled (**Fig. S8**). Logically, the bias of heterochromatic chrY regions for smaller fragments was also abolished in cancer (**Fig. 3E** to compare with **Fig. 1I**). In our local dataset, the same pattern was observed though depletion of fragments below 147 bp was comparable in magnitude between colorectal cancer patients and healthy individuals (**Fig. S9**).

In the Cristiano cohort, we classified fragments as short (100–147 bp) or long (147–200 bp). Fragments below 100 bp were excluded from the short-fragment category due to their very low frequency in Cristiano data. The 147 bp and 200 bp boundaries were chosen as the lengths at which the length relative frequency difference between chrY and auto+X was null. In male healthy individuals, the proportion of short fragments was higher in chrY than in auto+X, while the proportion of long fragments was lower (**Fig. 3F**). In male cancer patients, this pattern persisted, but it was much attenuated, particularly for short fragments (**Fig. 3G**), consistent with our earlier observations.

### ChrY fragmentation as a cancer biomarker

We showed above that chrY and auto+X exhibits distinct fragmentation profiles in healthy individuals, and this distinction is attenuated in cancer patients. We therefore thought that chrY fragmentation could serve as a potential biomarker for cancer detection in male patients. We computed long- and short-fragment proportion ratios for all male individuals in the Cristiano dataset (Materials and Methods). A two-dimensional representation of these two ratios showed a clear separation between the healthy and cancer data sets, suggesting that healthy and cancer individuals could be discriminated based on these ratios (**Fig. 4A**).

**Figure 4.**
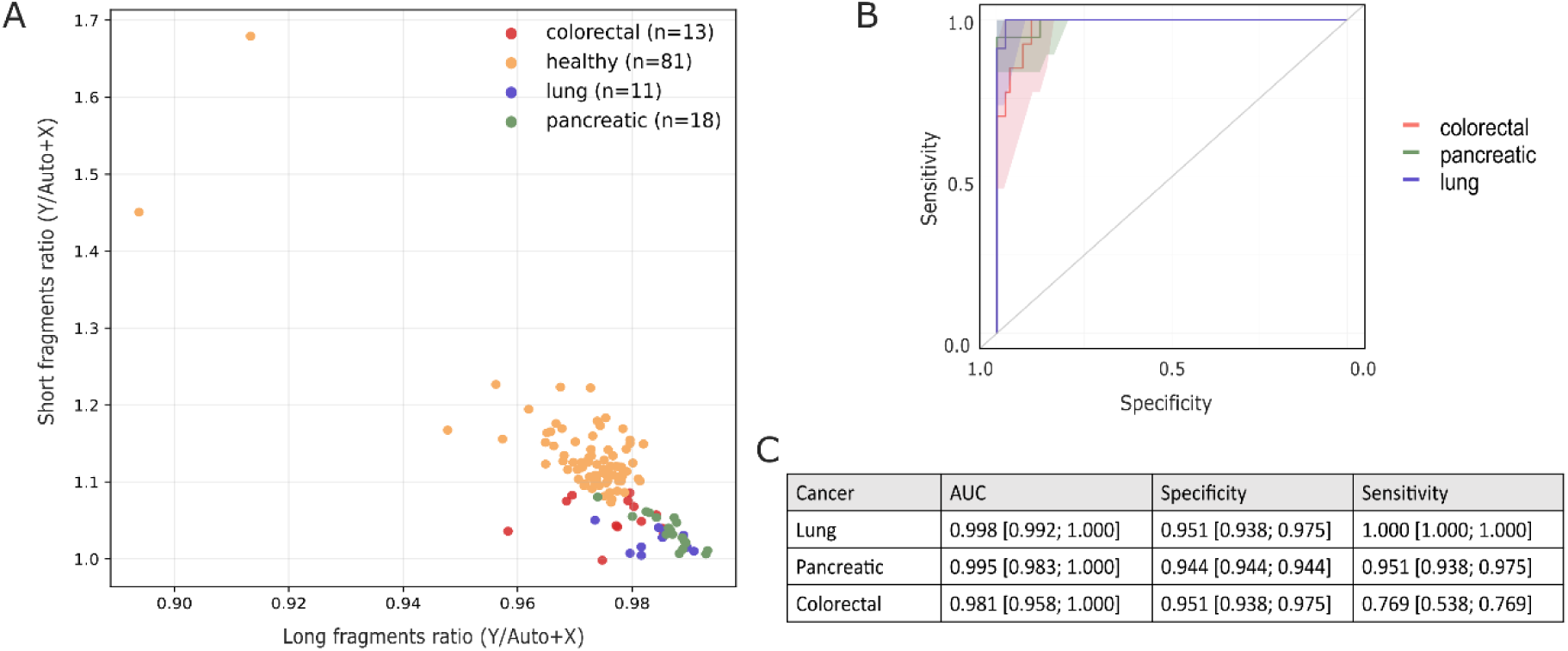
(**A**) 2D representation of short (100 – 147 bp) *versus* long (147 – 200 bp) fragment proportion ratios for male healthy, colorectal, lung and pancreatic cancer patients (Cristiano cohorts). (**B**) Receiver operating characteristics curve representing the performance of the SVM model to discriminate male colorectal, pancreatic and lung cancer patients from male healthy individuals (Cristiano cohorts). (**C**) Performance metrics of the SVM model to discriminate male colorectal, pancreatic and lung cancer patients versus male healthy individuals (Cristiano cohort).

We trained a Support Vector Machine (SVM) model (26) using the long- and short-fragment proportion ratios as input features. Leave-one-out cross-validation was employed for performance assessment given the limited number of cancer samples. The model discriminated colorectal, pancreatic, and lung cancer patients from healthy individuals with high performance, achieving sensitivities of 0.769, 0.951, and 1.00, respectively, at specificities close to 0.95 (**Figs 4BC**).

## DISCUSSION

In this study, we asked whether specific chromosomes display atypical fragmentation patterns in healthy individuals and how these patterns are altered in cancer. Using Cristiano *et al.* cohorts (5), we systematically compared the fragment length distribution of each chromosome with that of all the other chromosomes using Kullback-Leibler divergence. In healthy males (n = 81), chrY showed the greatest divergence, as expected from our previous observations (17), followed by chr19, chr17 and chr22. The latter three chromosomes were not anticipated to display such distinct fragmentation profiles, but are among the chromosomes with the highest gene density (27). Notably, chrY was enriched in cirDNA fragments shorter than 147 bp, whereas chr19, chr17, and chr22 showed enrichment in both the shorter and the longer ends of the fragment length distribution (> 167 bp), indicating distinct modes of divergence from the genome-wide pattern. The enrichment for shorter fragments among chrY-derived cirDNA compared to autosomes and chrX-derived cirDNA was reproducible in a local healthy cohort, although the transition from depletion to enrichment was close to 167 bp instead of being close to 147 bp. We interpret this difference as potentially resulting from our data to be much richer in cirDNA fragments < 100 bp compared to the Cristiano dataset, but our local dataset was small. This hence suggests that the location of the transition between depletion and enrichment might need to be adapted to each experimental protocol. Analysis of healthy females of the Cristiano dataset (n = 164) identified the same three autosomes as divergent, with comparable fragmentation profiles. To include chrY in our analyses, we focused on male individuals in subsequent analyses.

Further analysis revealed that the excess of short chrY-derived fragments predominantly originated from heterochromatic regions. This finding was unexpected because heterochromatin is generally associated with histone H1 enrichment and compact chromatin, which is expected to reduce accessibility to nucleases (28,29). The distinct behavior of chrY may reflect its unusual genomic and nuclear organization. ChrY contains the largest constitutive heterochromatic region in the human genome, and its size varies substantially among individuals (from 45 to 85 Mbp), with much of this variation attributable to heterochromatin (14). In addition, chrY is the smallest and least gene-dense human chromosome, and chromosome size and gene density have been linked to nuclear spatial organization (30,31). These properties, together with reported effects of chromosome organization on micronucleus stability (31), may contribute to a distinct chromatin environment before the release of chrY-derived DNA into the circulation. The precise mechanisms underlying the preferential generation of short fragments from chrY heterochromatin, however, remain to be established.

In contrast, the three divergent autosomes chr19, chr17, and chr22 displayed associations between short cirDNA fragments and transcriptionally active genomic features, including TSS, promoters and DNase hypersensitive regions (32). This is consistent with their high gene density, which increases the relative representation of such regulatory regions. Longer cirDNA fragments were additionally associated with heterochromatic regions of these chromosomes. Together, these observations suggest that chromosome-specific fragment length distributions can arise from distinct combinations of genomic composition and chromatin organization. It should also be noted that the Cristiano dataset contains very few fragments shorter than 100 bp. Consequently, additional chromosome- or region-specific fragmentation patterns may remain undetected at shorter lengths.

Because cirDNA contains information about nucleosome positioning (10,33,34), we next compared nucleosome occupancy across chromosomes in healthy individuals. Autosomes and chrX showed broadly comparable nucleosome densities, whereas chrY displayed at least threefold lower nucleosome occupancy. This observation was independently supported by MNase digestion data from a hematopoietic cell line. The reduced nucleosome occupancy of chrY was unexpected given its extensive heterochromatic content, as heterochromatin is frequently associated with high nucleosome density (35). It is unlikely that the bias towards shorter chrY-derived cirDNA fragment lengths is related to nucleosome density since nucleosome depletion usually results in the complete degradation of DNA by the DNAses (36) and the corresponding cirDNA is not observable. Moreover, we did not detect a significant association between nucleosome density and specific chrY chromatin regions. These results thus suggest that the environment and potentially the management of chrY is different, and chrY-derived cirDNA properties reflect these differences.

Potential mapping biases should also be considered. Although PAR were excluded and low-quality reads were filtered, the presence of the X-transposed region and the abundance of repetitive sequences in chrY heterochromatin could affect read mapping and therefore fragment quantification (37). Such biases could particularly lead to underrepresentation of fragments originating from repetitive regions. Nevertheless, the independent MNase-based observation of reduced chrY nucleosome occupancy supports the presence of genuine structural or epigenetic differences associated with chrY, although further experimental validation will be required to establish their molecular basis.

Since cancer is known to induce changes in cirDNA fragmentation profiles (4,11), we next investigated whether the chromosome-specific features above were altered in cancer. We found no observable difference regarding chrY nucleosome density. However, our approach could not detect chromatin remodeling or shifts in nucleosome positioning, both of which are known to occur in cancer (38,39). We also found no change in the fragmentation of chr19, chr17, and chr22. In striking contrast, the characteristic chrY fragmentation pattern was substantially attenuated. Specifically, the excess of short chrY-derived fragments observed in healthy males was markedly reduced in cancer, such that chrY fragment lengths no longer showed the same degree of deviation from the genome-wide distribution. This effect was particularly evident in the heterochromatic regions that contributed predominantly to the short-fragment population in healthy individuals.

The remarkable behavior of chrY-derived cirDNA fragment logically suggested to use the ratios of short and long cirDNA fragments between chrY and all the other chromosomes as a cancer biomarker for male patients. We showed that a ML model could indeed exploit chrY cancer variation to discriminate healthy males from patients with colorectal, pancreatic and lung cancers, with AUCs > 0.98. These results indicate that chromosome-specific fragmentation features can provide information that may be obscured when fragment-length distributions are considered at the genome-wide level.

Our findings also relate to previous observations linking chrY to cancer biology through mosaic loss of chrY (mLOY) (40,41). Although mLOY and the fragmentation phenotype described here represent distinct phenomena, both highlight the potential relevance of chrY-specific alterations in cancer. Our results also relate to the DELFI approach, which uses genome-wide fragment-length distributions within subchromosomal windows and ML for cancer detection (5). In males, incorporating chrY-specific fragmentation features into such frameworks could provide an additional source of information, particularly within chrY heterochromatic regions.

Overall, this study identifies an unusual fragmentation and nucleosome-occupancy profile associated with chrY-derived cirDNA. The preferential generation of short fragments from chrY heterochromatin, together with the markedly reduced nucleosome occupancy of chrY, points to distinctive structural and epigenetic properties of this chromosome. The attenuation of this fragmentation signature in cancer further suggests that chromosome-specific cirDNA fragmentation can capture cancer-associated alterations that are not apparent from genome-wide analyses. Determining the molecular mechanisms underlying these observations, and assessing their reproducibility across larger and clinically diverse cohorts, will be important for establishing the biological and translational significance of chrY-derived cirDNA fragmentation.

## MATERIAL & METHODS

### Healthy individuals and patients

We downloaded cirDNA fragment data for healthy, colorectal cancer, lung cancer, pancreatic cancer and prostate cancer samples corresponding to the Cristiano, *et al.* (5) study, and the Adalsteinsson, *et al.* (25) study, from FinaleDB (42). For lung cancer, samples corresponding to post-treatment conditions we excluded, and for healthy individuals, technical replicates were excluded. For each dataset, we only kept samples from male individuals: 81 healthy, 13 colorectal cancer, 11 lung cancer, and 18 pancreatic for Cristiano, and 557 prostate cancer for Adalsteinsson.

The local cohort comprised 20 healthy male samples and 8 stage IV metastatic colorectal cancer samples from the screening procedure of the UCGI 28 PANIRINOX study (NCT02980510/ EUCT 2024-510645-34-00). Written informed consent was obtained from all participants in the local cohorts. All protocols and patients information are available from Piserva, *et al*. (43).

### Preprocessing

For the Cristiano and Adalsteinsson cohorts, files downloaded from FinaleDB contain the position of cirDNA fragments mapped on the human genome GRCh38 in a BED-like tab-separated value format. Fragments with a quality score < 30 were filtered out. BED-like files were converted into genuine BED format using the BEDTools suite. For the local cohort, we converted BAM files to BED format using the BEDTools suite.

For all datasets, fragments mapped on chrY were extracted in separate BED files. Fragments mapped in PAR regions of chrX were discarded, in order to avoid biases due to homologous sequences. In chrY, three regions that did not contain any fragments, including the PAR regions, were discarded.

### Fragment length analysis

For each dataset, fragment length distributions were computed separately for the autosomes + chrX (auto+X) origin and for the chrY origin. The distributions were estimated by the relative frequencies of each fragment length. In order to compare fragmentation profiles between chrY and auto+X, we determined the difference in relative frequencies (ΔF) at each fragment length. Repeated random subsampling was applied to select an equivalent number of fragments from the auto+X and chrY conditions, and bootstrapping was used to estimate confidence intervals.

Long and short fragment proportion ratios were computed for all the samples as follows. The proportion of long (147 to 200 bp) and short fragments (100 to 147 bp) was computed for chrY and auto+X and named 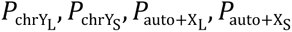, respectively. Ratios of these proportions were then computed for each sample:

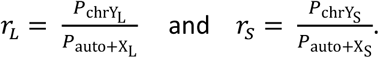

### Nucleosome occupancy analysis

Nucleosome positions were inferred separately for the Cristiano healthy and Adalsteinsson datasets using the algorithm described in Ref. (11). Nucleosome positions were then counted for each chromosome *per* 10kb genomic bin. Centromeric regions were excluded.

### MNase data analysis

We downloaded public data of MNase-digested chromatin from a cell line of activated human CD4+ T cells (23). We considered that the fragments obtained by MNase digestion were attached to nucleosomes, and counted the fragments for each chromosome *per* 10 kb genomic bin.

### Machine learning

Machine learning was performed using R (v 4.3.3) and the software packages: caret (v 7.0-1) for building the model for cross-validation, and pROC (v 1.18.5) for AUC calculation. We performed leave-one-out cross validation using a SVM model. The imbalance of the healthy and cancer classes distribution was corrected using SMOTE.

## DECLARATIONS

## Ethics approval and consent to participate

Written informed consent was obtained from all participants in the local cohorts. The study was approved by local ethics committees and conducted in accordance with the Declaration of Helsinki.

## Availability of data and material

The Cristiano and Adalsteinsson datasets used in this study are publicly available from FinaleDB (42) at https://finaledb.research.cchmc.org/query. The data from the local cohort can be retrieved from the European Genome-phenome Archive with accession number EGAS50000001319.

All the computer codes developed to perform the analysis are available from GitHub at https://github.com/mrichaud/chrY_cirDNA.git

## Authors’ contributions

L.R. and M.R. did the code and analysis and prepared all figures. M.R. and J.C. wrote the main manuscript text. E.P., J.C. and A.R.T. corrected and improved the manuscript text and figures. J.C. and A.R.T. conceptualized the study and provided funding.

