## Supplemental information for "Distinctive fragmentation of circulating DNA from chromosome Y and its alteration in cancer"

### SUPPLEMENTARY FIGURES

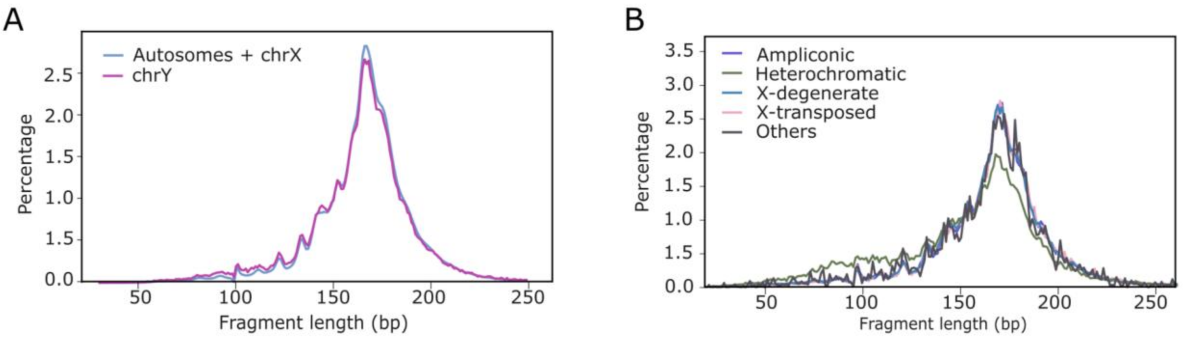


**Figure S1.** Length distribution of healthy individuals from the local cohort of chrY-derived fragments *versus* autosomes and chrX-derived fragments (**A**), and of chrY-derived fragments for each subregion type (**B**).

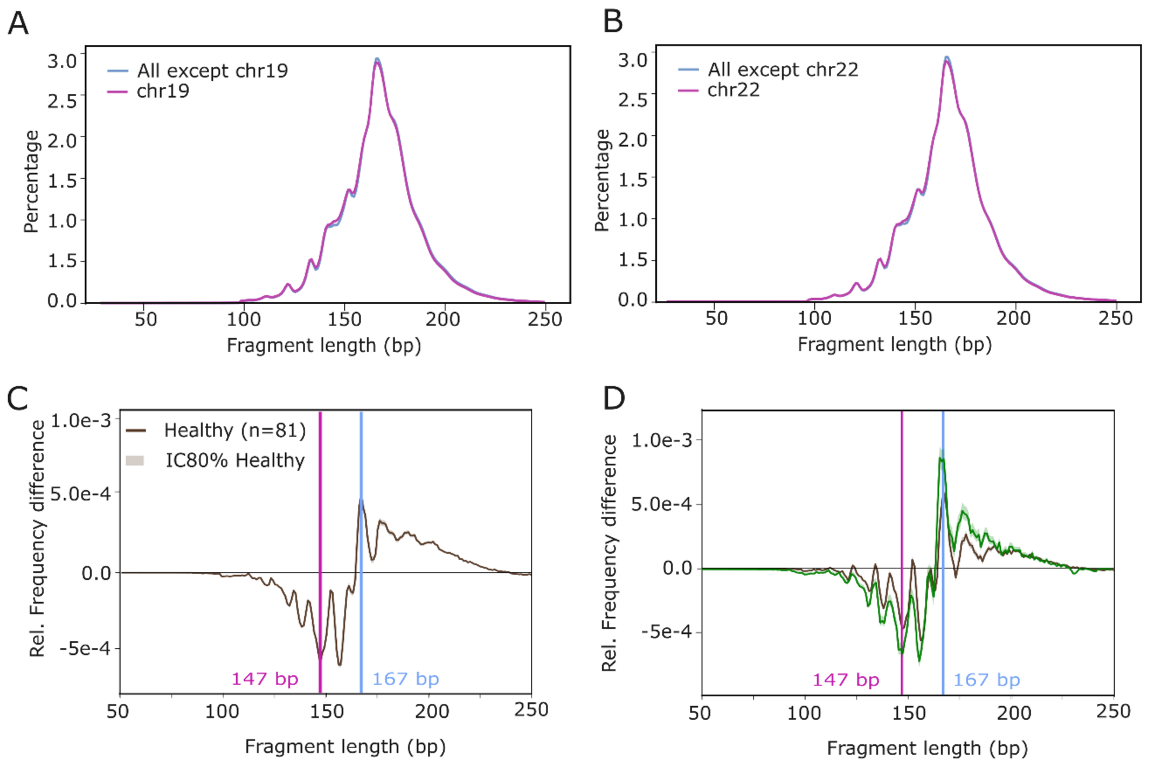


**Figure S2.** Fragment length distribution divergence of chr17 and chr22 (**A, B**), also represented as relative frequency differences (**C, D**).

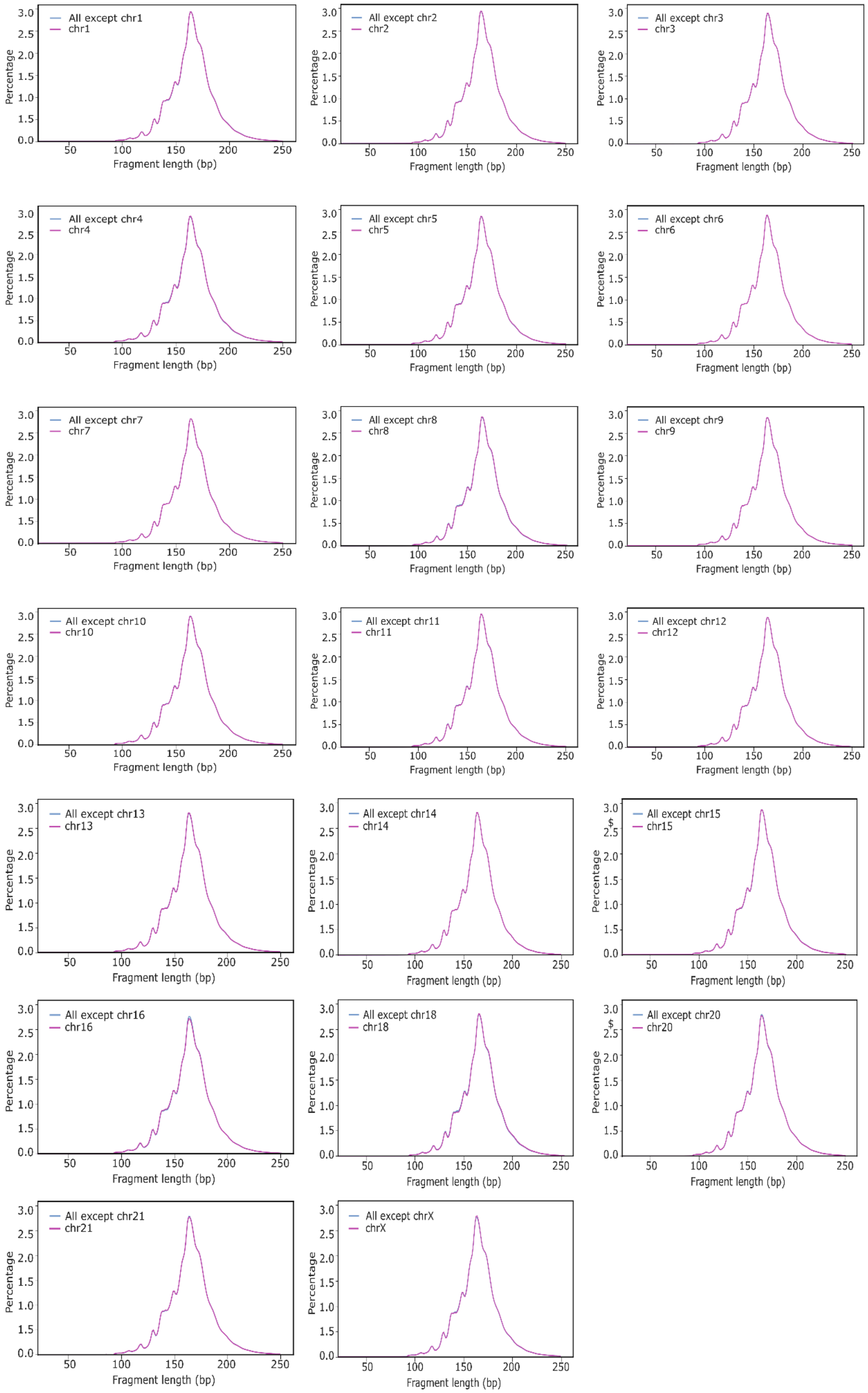


**Figure S3.** Fragment length distributions of non-divergent chromosomes.

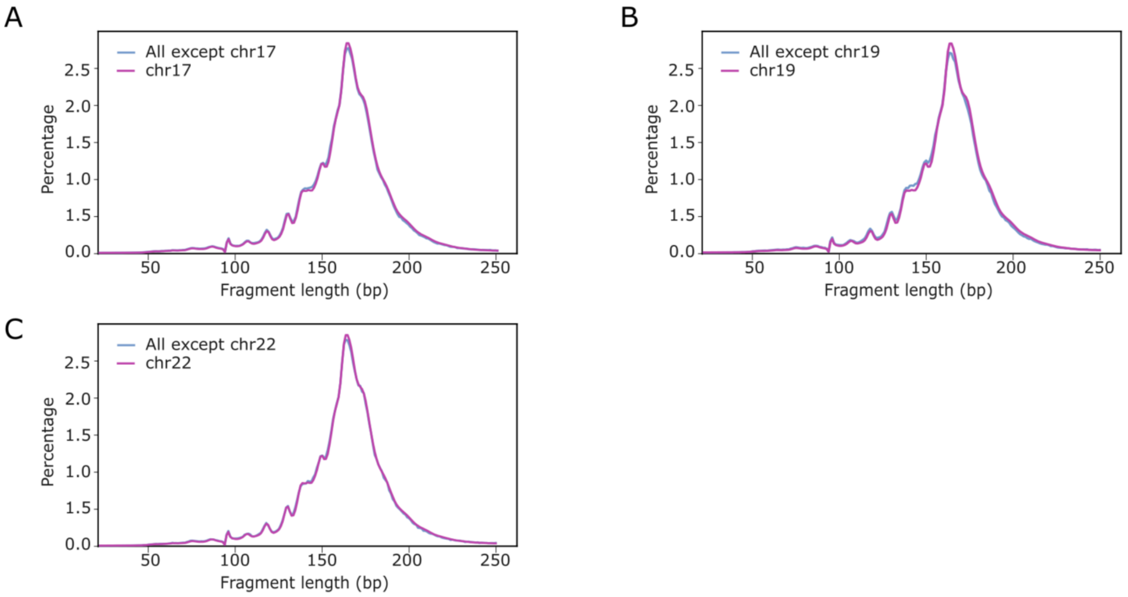


**Figure S4.** Fragment length distributions of chr17 (**A**), chr19 (**B**) and chr22 (**C**) for healthy male samples of the local cohort.

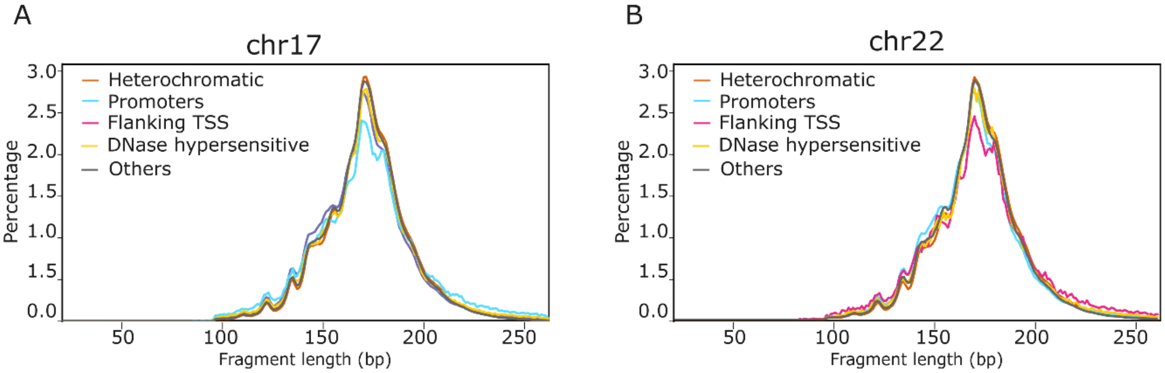


**Figure S5.** Association of fragment lengths and genomic regions for chr17 (**A**) and chr22 (**B**).

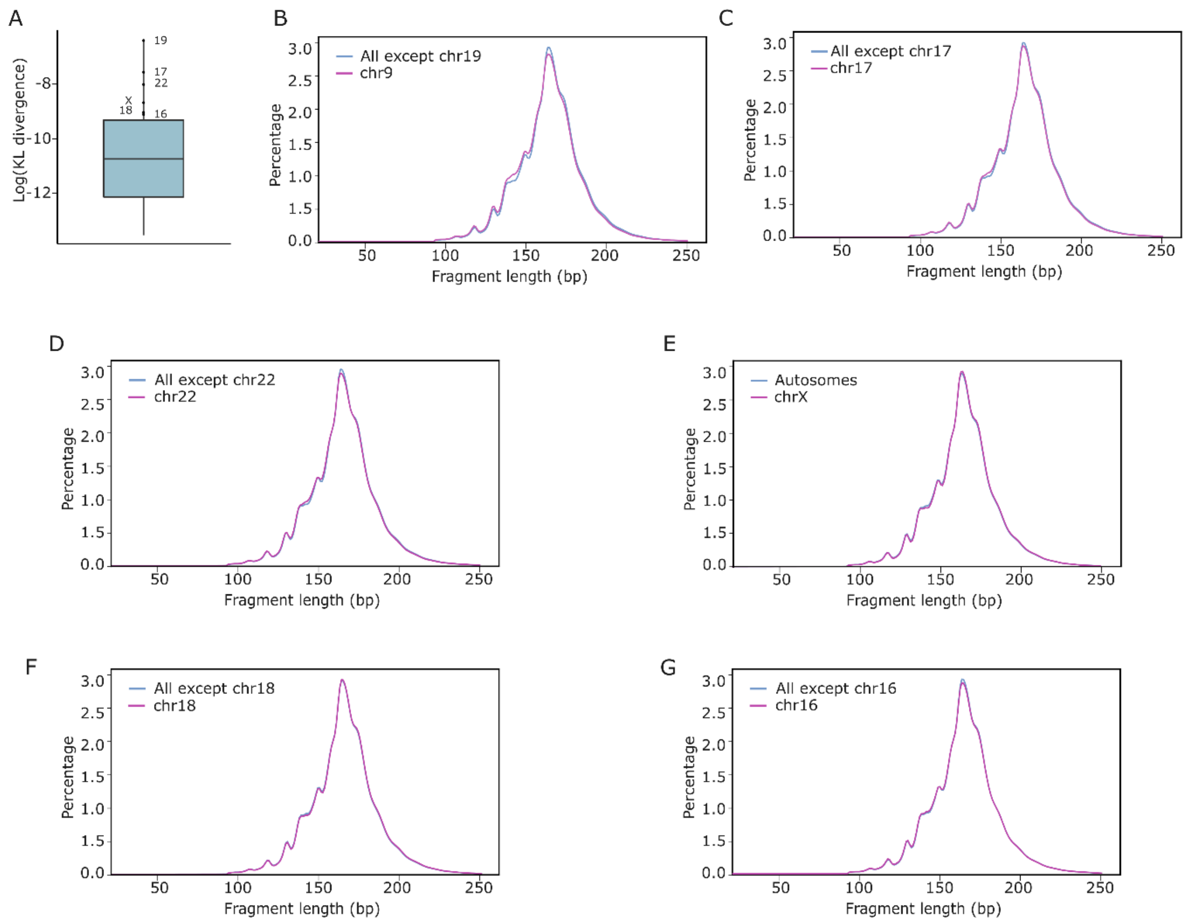


**Figure S6.** Fragment length distribution divergence in female healthy individuals (Cristiano cohort). The same autosomes are most divergent (**A**) with additional, marginal divergence of chrX, chr18 and chr16 (**B-G**).

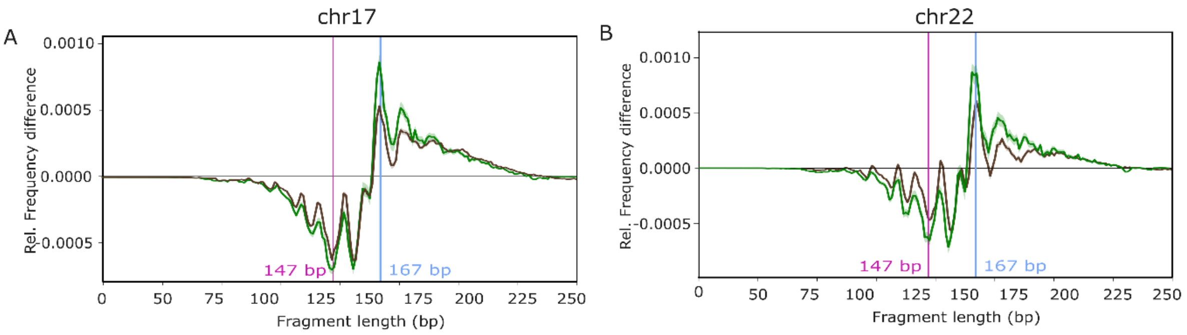


**Figure S7.** CirDNA fragment length relative frequency differences between chr17 and all other chromosomes in healthy individual (black) and pancreatic cancer (green) (**A**). (**B**) Same for chr22.

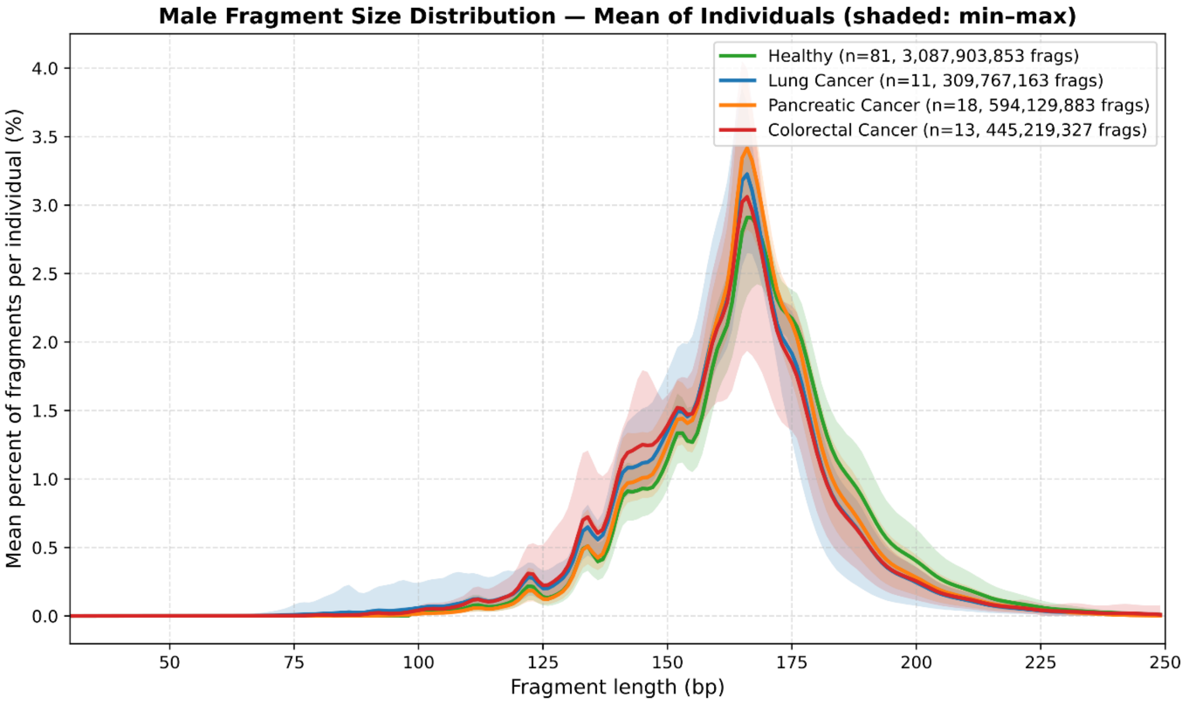


**Figure S8.** Illustration in Cristiano data of the classical phenomenon of shorter cirDNA fragment enrichment in cancer cohorts compared to healthy, colorectal being often the most pronounced.

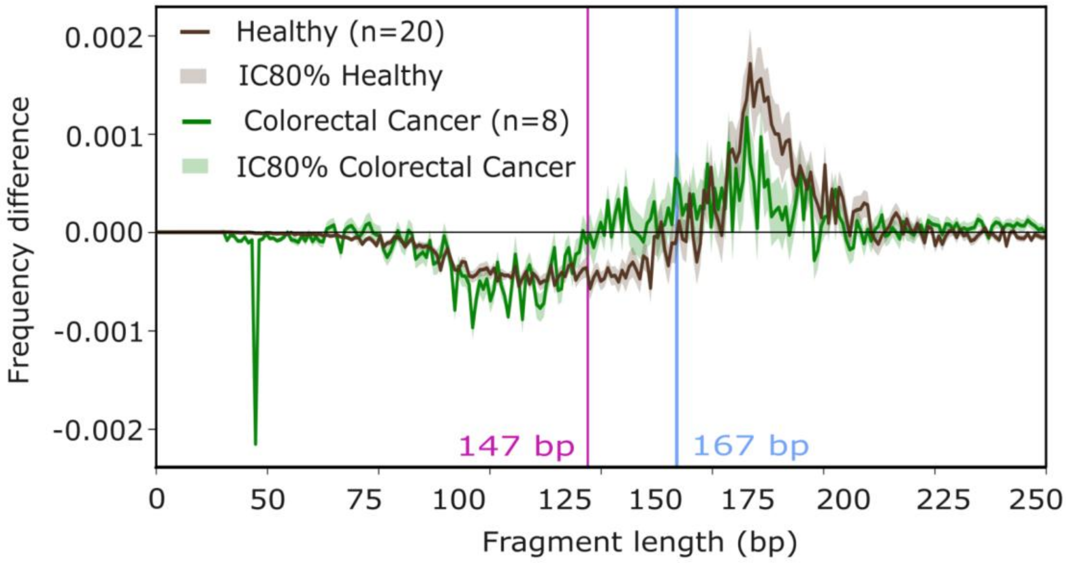


**Figure S9.** Fragment length relative frequency differences between chrY and autosomes+chrX in male healthy individuals and colorectal cancer patients (local cohort).
